# In Vitro Safety “Clinical Trial” of the Cardiac Liability of Hydroxychloroquine and Azithromycin as COVID19 Polytherapy

**DOI:** 10.1101/2020.12.21.423869

**Authors:** Bérénice Charrez, Verena Charwat, Brian Siemons, Henrik Finsberg, Evan Miller, Andrew G. Edwards, Kevin E. Healy

## Abstract

Despite global efforts, there are no effective FDA-approved medicines for the treatment of SARS-CoV-2 infection. Potential therapeutics focus on repurposed drugs, some with cardiac liabilities. Here we report on a preclinical drug screening platform, a cardiac microphysiological system (MPS), to assess cardiotoxicity associated with hydroxychloroquine (HCQ) and azithromycin (AZM) polytherapy in a mock clinical trial. The MPS contained human heart muscle derived from patient-specific induced pluripotent stem cells. The effect of drug response was measured using outputs that correlate with clinical measurements such as QT interval (action potential duration) and drug-biomarker pairing.

Chronic exposure to HCQ alone elicited early afterdepolarizations (EADs) and increased QT interval from day 6 onwards. AZM alone elicited an increase in QT interval from day 7 onwards and arrhythmias were observed at days 8 and 10. Monotherapy results closely mimicked clinical trial outcomes. Upon chronic exposure to HCQ and AZM polytherapy, we observed an increase in QT interval on days 4-8.. Interestingly, a decrease in arrhythmias and instabilities was observed in polytherapy relative to monotherapy, in concordance with published clinical trials. Furthermore, biomarkers, most of them measurable in patients’ serum, were identified for negative effects of single drug or polytherapy on tissue contractile function, morphology, and antioxidant protection.

The cardiac MPS can predict clinical arrhythmias associated with QT prolongation and rhythm instabilities. This high content system can help clinicians design their trials, rapidly project cardiac outcomes, and define new monitoring biomarkers to accelerate access of patients to safe COVID-19 therapeutics.

## Introduction

When the World Health Organization declared a global pandemic on March 11th 2020, little was known about the pathogenesis of the severe acute respiratory syndrome coronavirus 2 (SARS-CoV-2). It was described and treated as a respiratory disease, in which the virus targeted the epithelial cells of the respiratory tract, resulting in alveolar damage, edema and fibrosis. Now, with more than 45 million cases and a million deaths worldwide, there is clinical evidence that the virus also has non-negligible long-term effects on multiple organs, including heart, kidney, vasculature, liver and even brain[1–7].

With the absence of FDA-approved medicines for the treatment or prevention of COVID-19, clinicians have been pressed to treat patients in critical stages without FDA approved protocols. They have therefore relied on several small scale clinical studies to repurpose compounds approved by regulatory bodies as monotherapies in the hope of improving patient outcomes. Early clinical trials identified chloroquine (**CQ**), hydroxychloroquine (**HCQ**) and azithromycin (**AZM**) as promising drugs to help treat or reduce the effects of SARS-CoV-2 [8–12]. A non-randomized clinical trial in France identified HCQ, in combination with AZM, as being capable of significantly reducing respiratory viral loads [8]. This was confirmed by more recent retrospective studies[13, 14], but was also heavily criticized and refuted by other recent studies employing *a priori* designs[15–18]. Several additional small clinical trials have shown mixed outcomes for HCQ treatment of COVID-19 patients[19].

A serious concern with these studies is that patients were treated with drugs that have known cardiac complications, and their effects on the heart in polytherapy were unknown. HCQ inhibits *hERG* (***I***_***Kr***_) potassium channels, it is known to increase in QT interval of cardiomyocytes, and can induce arrhythmias that are responsible for sudden death [20]. AZM is also associated with an increased risk of cardiovascular death, due to Torsade de Pointes (**TdP**) and polymorphic ventricular tachycardia [21]. With respect to polytherapy, clinical trials have demonstrated a synergistic effect of HCQ and AZM to prolong QT interval [22, 23]; however, alterations in arrhythmic event frequency were controversial when compared to HCQ or AZM alone. In the absence of rapid clinical trials for polytherapy safety, there is an urgent need for screening tools to increase the speed at which potential therapeutics are evaluated for cardiac liabilities. Common *in vitro* systems used for cardiac drug screening include cell 2D monolayers and animal testing, which often fail to replicate human physiology, particularly electrophysiology, and pharmacokinetic properties [24]. Engineered heart tissue, organoids or microfluidic-based microphysiological system are emerging alternatives for state-of-the-art drug screening [25, 26]. With the rapid spread of COVID-19, microphysiological systems (**MPS**) have recently shown to be a promising tool to study virus entry and replication mechanisms, subsequent cytokine production, as well as effects of existing and novel therapeutics or vaccines [27, 28].

In this paper, we demonstrate the utility of a cardiac MPS (**Figure 1**) for determining the cardiac liability associated with HCQ and AZM polytherapy in an *in vitro* design analogous to a Phase I safety clinical trial. Our cardiac MPS contains a three-dimensional (**3D**) cell chamber in which human induced pluripotent stem cell-derived cardiomyocytes (**hiPSC-CMs**) are confined and self-assemble to form uniaxially beating heart muscle [24, 29]. HiPSC-CMs have been successfully used for *in vitro* assessment of drug-induced arrhythmias, especially since they respond consistently to *hERG* channel block (QT prolongation and arrhythmias) and calcium channel block (action potential duration shortening, impaired contractile function) [6, 30]. It makes them excellent candidates to screen for cardiac liability of HCQ and AZM, both of which are known to block hERG channels, and also act upon other cardiac ion channels[31]. By assaying hiPSC-CMs expressing a genetically encoded calcium sensor (GcAMP6f), loaded with a voltage-sensitive fluorescent probe (BeRST), we assessed the electrophysiology and calcium handling of the tissues during serial drug exposures. We showed that HCQ and AZM significantly increase 80% repolarization time (APD_80_) and rhythm instabilities, starting at clinically relevant exposure days, and were accompanied with EAD and TdP instances. HCQ+AZM combination also showed a significant increase in APD_80_, however, few instabilities or arrhythmic events were observed. Finally, proteomics analysis of cell culture effluent enabled detection of biomarkers that were directly correlated with cardiotoxicity, apoptosis and contraction mechanics alteration.

**Figure 1.**
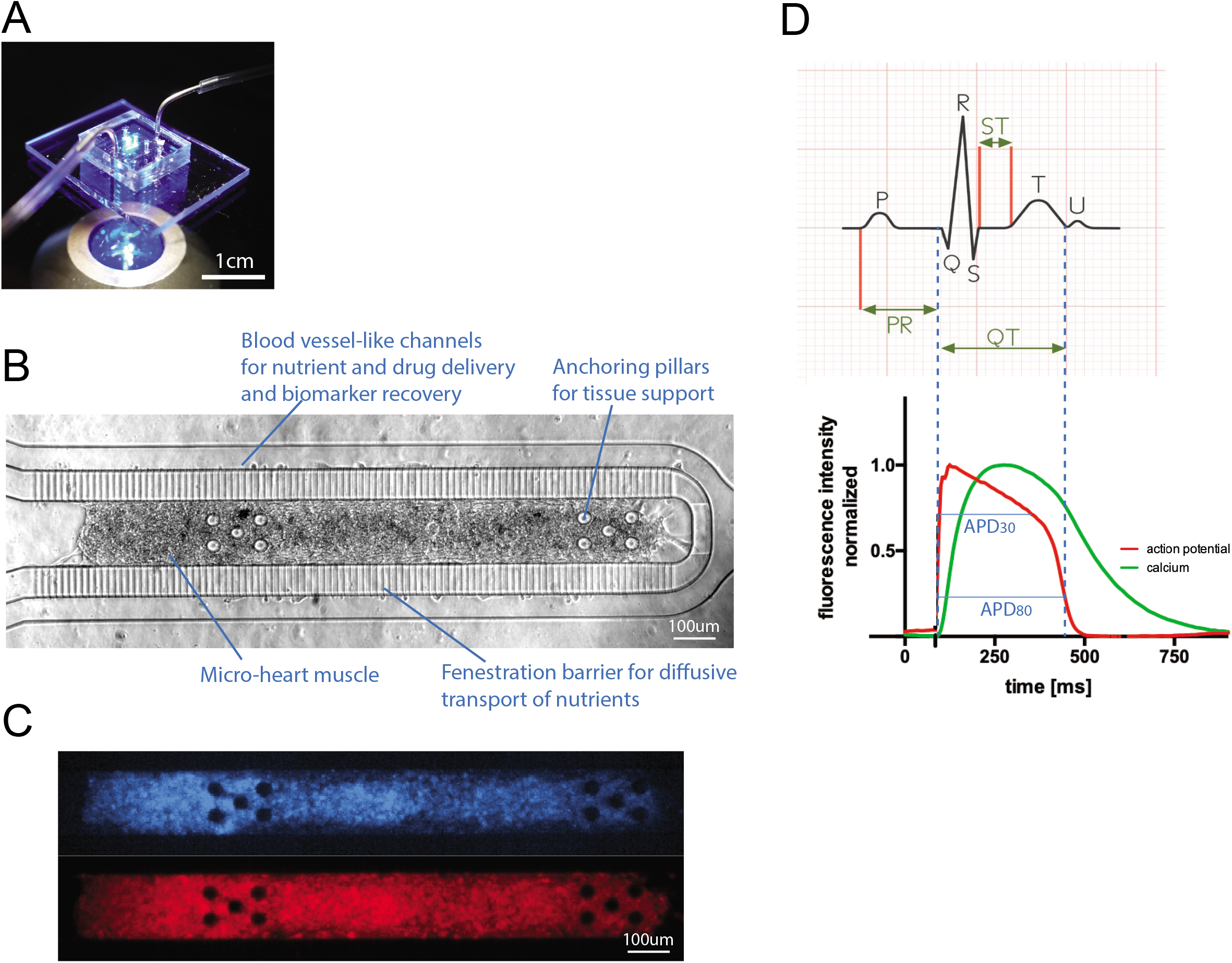
The cardiac microphysiological system. (**A**) Photograph of a cardiac MPS in fluorescent light with feeding tubing. **(B)** Brightfield image of a cardiac MPS loaded with 4,000 human induced pluripotent stem cell-derived cardiomyocytes. The cell chamber is separated from adjacent feeding channels via a fenestration barrier of 2μm wide grooves allowing for nutrient diffusion while protecting the tissue in the cell chamber from media flow-induced shear stress. The anchoring pillars on either side of the cell chamber help keep the heart muscle elongated and provide resistance for contraction. (**C**) Representative images of the same tissue under GFP fluorescence for calcium transient recordings (top) or FarRed voltage dye staining (bottom). (**D**) The top graph shows a typical ECG recording from which the clinical QT interval can be determined. We use APD_80_ as a proxy for QT duration, corresponding to the duration of the action potential at 80% of its repolarization (bottom). The APD (red) and Ca (green) waveforms are timestamped identifying temporal kinetics.

## Results and Discussion

### Chronic exposure to HCQ for 10 days resulted in QT prolongation and rhythm instabilities that correlated with arrhythmic events and clinical observations

Chronic exposure to an HCQ dose mimicking clinical protocols decreased the beat rate starting at day 4 with a statistically signifcant decrease on days 4, 7 and 8 (**Figure 2A**). APD_80_ increased markedly from day 7 onwards (**Figure 2B**), with the maximum APD_80_ increase reaching 850ms. All but one tissue exhibited an APD_80_ at least 400 ms longer on day 7 and day 9 than prior to HCQ exposure. These arrhythmogenic changes to the AP were tightly correlated with the appearance of instabilities in the HCQ tissues from day 5 onwards (**Figure 3A**). APD_80_ was directly correlated to calcium transient duration (CaD) (**Supp. Figure 1D**), and an increase of CaD_80_ above 600ms was observed at day 2 and increased over the duration of HCQ treatment. Arrhythmic events began appearing at day 4 in 50% of the tissues. At day 7, tissues exhibited both arrhythmogenic AP waveforms and CaD_80_ increase, and by days 9 and 10, 50% of tissues exhibited weak or no dynamic signal change, suggesting loss of resting membrane potential (**Figure 3D**). Representative traces of arrhythmic events are shown in **Figure 3 G-J**, comparing 30 second calcium traces at day 0 and day 9 (**Figure 3G**). Late calcium peaks are a signature of EADs in membrane potential and were clearly observed (red arrows) in day 9 HCQ recordings, as were the marked increases in duration of the calcium transient, itself a marker for APD prolongation. Together these data reflect HCQ’s well known block of *I*_*Kr*_, which prolongs the QT interval and its *in vitro* proxy APD [22, 32] which are associated with arrhythmic events[20, 33, 34]. Our AP data, which is consistant with the clinical literature, indicates that the cardiac MPS system is a good predictor of clinical cardiotoxicity of HCQ.

**Figure 2.**
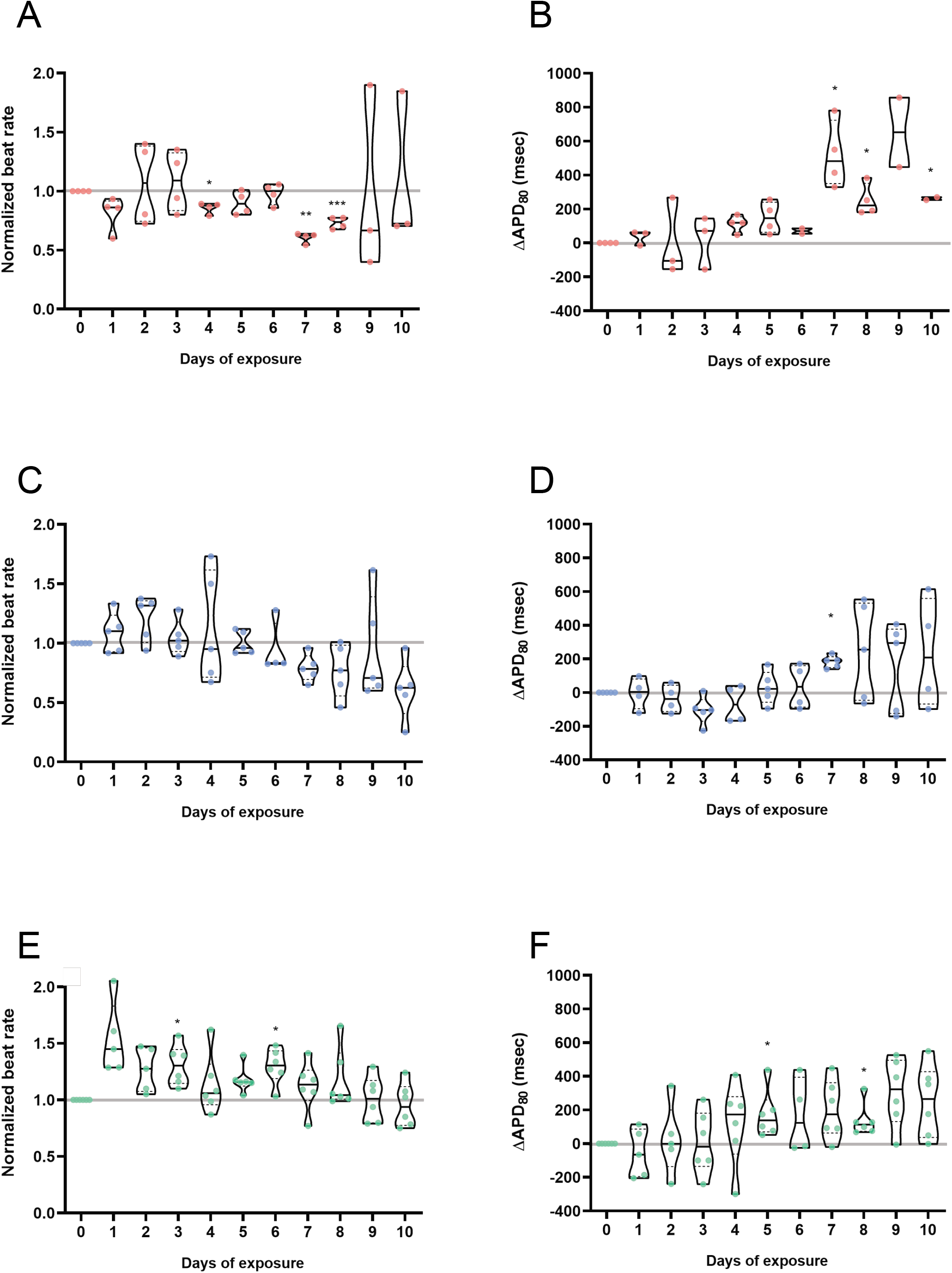
Electrophysiology analysis of chronic exposure to hydroxychloroquine (HCQ), azithromycin (AZM) or their polytherapy. Doses were chosen to closely mimic clinical trial drug prescription used for COVID-19 treatment: 0.24μM HCQ and 0.15 μM AZM on day1 followed by 0.12μM and 0.075 μM AZM on day 2 to day 10. Polytherapy was the combination of both monotherapy doses (**Supp. Table 1**). Violin plots demonstrate spontaneous beating during the therapy for HCQ (**A**), AZM(**C**) or polytherapy (**E**). Violin plots depicting the change in APD_80_ during the therapy for HCQ (**B**), AZM(D) or polytherapy (**F**). APD_80_ values were calculated from spontaneous recording, corrected for beat rate using the Fredericia correction[1]. In all graphs, the values were normalized to baseline at day 0 for each tissue. Each point corresponds to one heart muscle. The violin plots show the arithmetic median (solid line) and upper and lower quartile (dashed lines) as well as minimum and maximum values (truncation of violin shape). All tissues analyzed were within inclusion criteria of APD_80_ <500ms at baseline. Statistics run were one-way ANOVA repeated measures with multiple comparison to baseline day 0 and Dunnett’s post-hoc correction. * p<0.05; ** p<0.01; *** p<0.001.

**Figure 3.**
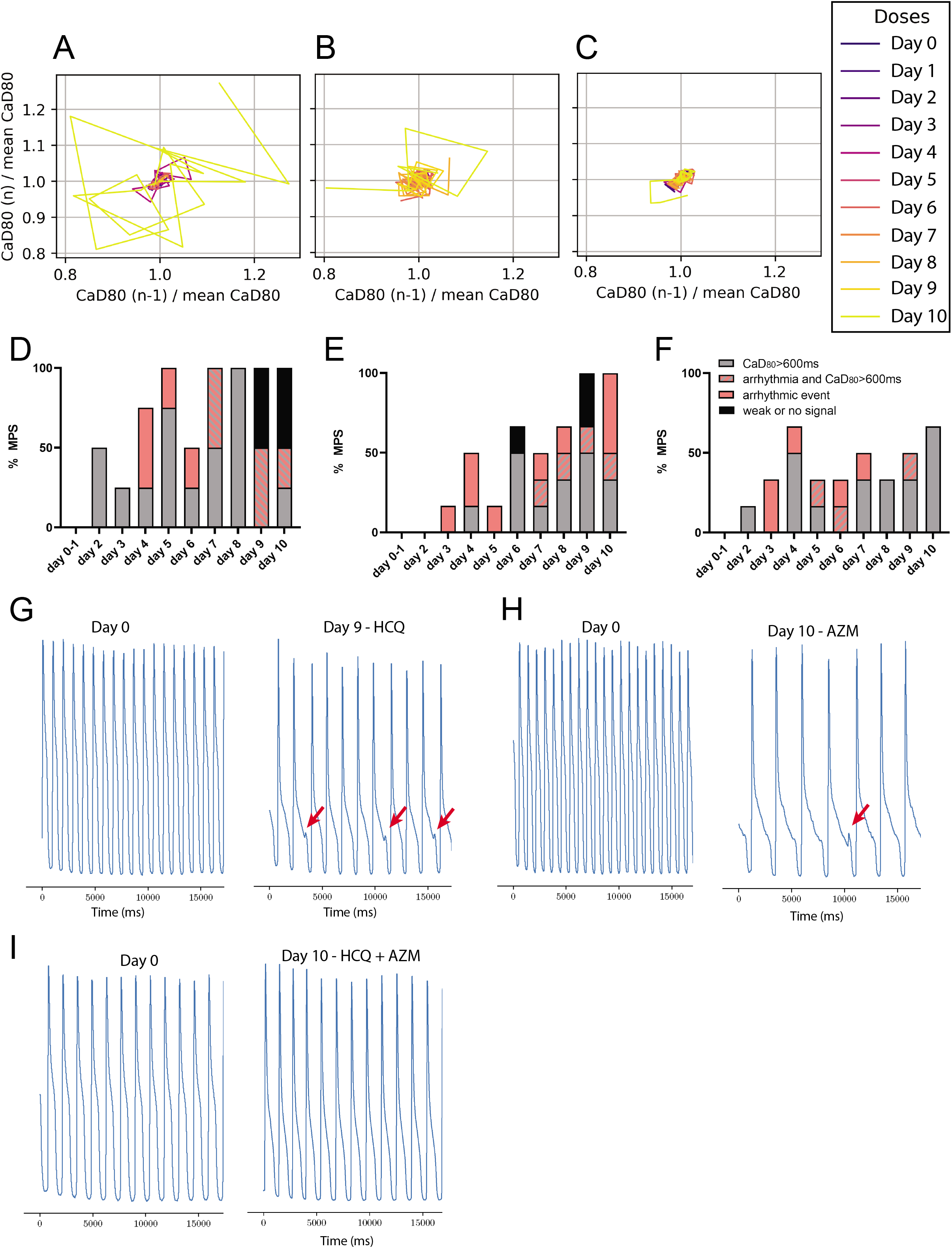
Instability and arrhythmic study of chronic exposure to hydroxychloroquine (HCQ), azithromycin (AZM) and their polytherapy. Poincare plots[2] were used to visualize rhythm instabilities in different tissues. Where small cluters of daily traces represent minimal arrhythmic risk, and large complex polygons are indicative of rhythm instability and high arrhythmic risk. **(A)** Representative Poincare graph of a tissue exposed to HCQ only. Disorganized polygons can be observed at day 5 and 10 indicative of drug-indueced arrhythmia. (**B**) Poincare graph of AZM-treated heart muscle where instabilities were observed starting on day 6. (**C**) Representative graph of polytherapy instabilities, which show fewer and smaller polygons when compared to monotherapy, indicative of reduced arrhythmic risk. (**D-F**) Histograms showing weak or no signal (black), CaD_80_ above 600ms with (striated) or without (grey) EAD and arrhythmic event (pink) as percentage of total heart muscles (i.e., MPS). The analysis was performed for HCQ alone (**D**), AZM alone (**E**), and for polytherapy (**F**). Representative calcium transient trace at baseline and after exposure to 9 days of HCQ (**G**) or 10 days of AZM **(H)** where EAD instances can be observed (red arrows). (**I**) Representative calcium transient trace at 10 day exposure polytherapy showing no EAD.

### Chronic exposure of AZM for 10 days showed QT prolongation and rhythm instabilities that correlated with arrhythmic events and clinical observations

Chronic AZM exposure did not significantly change beat rate (**Figure 2C**). However, as for HCQ, chronic AZM treatment increased APD_80_ by day 7, and a trend for persisting prolonged APD_80_ continued until day 10, albeit with large variation (**Figure 2D**). Triangulation also trended (p<0.2) to increase from day 8 onwards, although this did not reach significance (p=0.07 at day 10) (**Supp. Figure 1B**). Instabilities arose at day 6 and worsened over time (**Figure 3B**). Non-parametric analysis of instability showed a significant increase of chaotic polygons in AZM-treated tissues when compared with HCQ (p<0.05, not shown). CaD_80_ increased beyond 600ms at day 4, and day 6 onwards in 30-50% of the tissues. Arrhythmic events were observed from day 3 onwards, reaching 60% of tissues at day 10, with some tissues exhibiting both arrhythmia and CaD_80_ increase. Day 6 and 9 had respectively 16% and 33% of tissues with weak or no signal (**Figure 3E**). **Figure 3** shows a representative trace of arrhythmic events at day 0 versus day 10 (**Figure 3H**) of AZM exposure. Overall, APD_80_ increased on all treatment days compared to day 0, and EADs appeared after day 3, and were most prevalent on day 10. Since clinical AZM application is typically limited to 5 days, the observed incidence of pro-arrhythmic events during longer exposure times cannot be directly compared to clinical outcomes. However, AZM has been associated with increase in cardiovascular death, mostly through QT prolongation and arrhythmia [21, 35], and these outcomes are clearly indicated by our MPS measurements.

### Chronic exposure to both HCQ and AZM for 10 days showed QT prolongation and rhythm instabilities that correlated with arrhythic events and clinical observations

The beat rate increased at day 3 and 6; however, there was no clear overall trend (**Figure 2E**). APD_80_ was significantly increased on days 5 and 8, with a trend towards APD prolongation for all recordings after day 7 (p<0.2) (**Figure 2F**). This data set closely mimics clinical trials performed by Chorin et al. [36], where the QT interval increased starting at day 2 to day 5 with high variability in patient population. Triangulation was not significantly altered over the 10 days during our *in vitro* polytherapy trials, although there was a very slight trend towards increasing triangulation at day 4, 5 and 10 (p<0.2) (**Supp. Figure 1C**). Interestingly, instabilities were almost absent in this data set, when running non-parametric analysis on Poincare plots from the combination study (**Figure 3C**) versus those for individual HCQ (significant, p<0.05) or AZM (non significant trend) (**Figure 3 A,B)**. Although we observed a clear increase in APD_80_, and an increase of CaD_80_ above 600ms at day 2 and day 4-10 (**Figure 3E**), chronic polytherapy resulted in fewer arrhythmic events and only mild instability compared to monotherapy (**Figure 3I**). At most, 33% of tissues showed arrhythmia at day 3 and 6, with 16% arrhythmias at day 4, 5, 7 and 9 (**Figure 3F**). Non-parametric contigency analysis showed a decrease between EAD instances in polytherapy versus HCQ (significant, p<0.05) or AZM (trend, p<0.1) monotherapy. No tissues had weak signal or stopped beating. At the pathophysiologic level, this data fit well with prior studies describing the important role of AP triangulation in the transition from benign AP prolongation to unstable repolarization[37].

Together, these observations suggest that polytherapy rescues arrhythmogenesis resulting from the individual drugs. Recent clinical studies demonstrated chronic exposure to combination of HCQ and AZM led to QT increases with few arrhytmia events [17, 35, 38]. The concordance of the cardiac MPS data to arrive at similar conclusions demonstrates its power in predicting cardiac liabilities for combination therapy of repurposed drugs to treat SARS-CoV-2. Mechanisms explaining how arrhythmic events are absent despite a significant increase of the QT interval, can be complex and additional studies would be required to elucidate HCQ and AZM polytherapy-dependent mechanisms. However, based on the fact that HCQ and AZM have known multichannel blocking effects, and that *I*_*CaL*_ and *I*_*Na*_ block is known to reduce *I*_*Kr*_ dependent arrhythmias[33], we can hypothesize that the combination of both drugs can synergistically increase multichannel block responsible for lower arrhythmic instances when compared to monotherapies.

### Proteomics analysis of MPS effluent reveal candidate biomarkers for cardiotoxicity monitoring in patients treated with HCQ and AZM

For the polytherapy pharmacology study, media were analyzed for over 92 proteins as biomarkers of tissue injury (**Figure 4A**). The proteomics analysis of the MPS effluent included a wide array of biomarkers, most of them measurable in patients’ plasma, associated with different cardiac mechanisms, from morphology, cytoskeleton, mechanics to apoptosis and stress response. Cardiac troponin I (TNNI3), is a well known biomarker of cardiac injury and increased risk in mortality, common in COVID19 patients with underlying cardiovascular conditions[39]. Chronic exposure to the combination of AZM and HCQ showed no significant change in TNNI3 expression, suggesting that arrhythmic tissues are not undergoing major tissue damage. However, a clear decrease in erythropoietine (EPO) was observed. HPGDS, an intracellular enzyme that catalyzes the conversion of PGH2 to PGD2, was shown to decrease significantly. Interestingly, similar significant changes were also observed for carbonic anhydrase 14 (CA14) and tyrosine-protein kinase Fes/Fps (FES). These intracellular proteins are typically not secreted[40], and therefore are not strong biomarkers unless cells were damaged. The fact that alterations in levels for these proteins were observed in our study, suggests some degree of cell damage or stress, but not to the extent where troponin-actin complexes break down[41]. It is known that CA14 facilitates lactic acid transport across the cardiac sarcolemma [42], as well as improves myocardial energetics by facilitating mitochondria CO_2_ clearance [43]. We hypothesize the drug-related change in CA14 expression is a mechanism for the muscle to adapt antioxidant, contraction or waste management mechanisms to counter-balance cardiotoxic effects. Identification of biomarkers in the context of HCQ and AZM, or other polytherapies, will be a valuable tool in the design of COVID-19 therapeutics trials.

**Figure 4.**
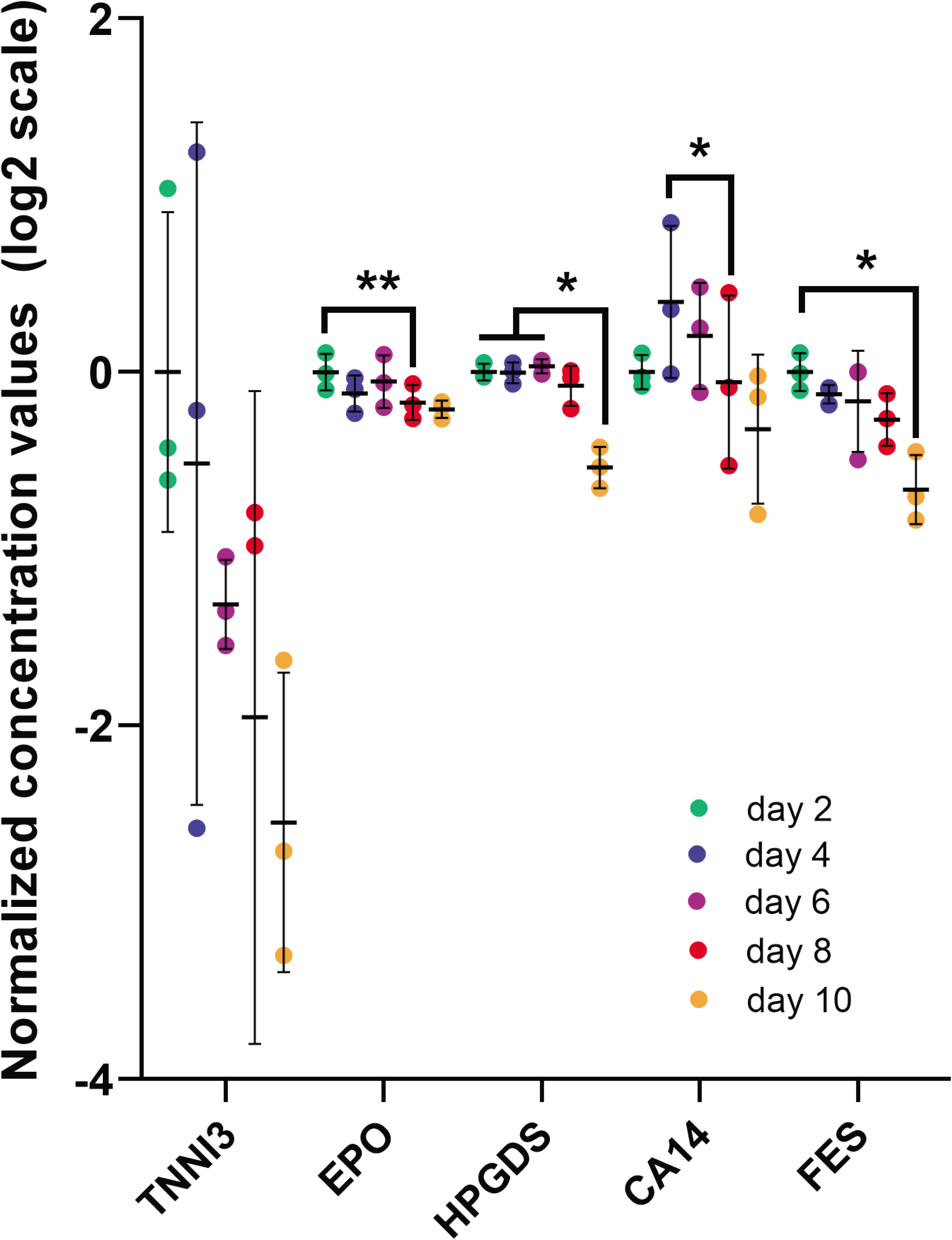
Proteomics analysis of microphysiological systems effluent during chronic exposure to HCQ and AZM polytherapy. Scatter plot of biomarkers showing significant changes with increasing doses of polytherapy in chronic study. The effluent media was analyzed on day 2, 4, 6, 8 and 10. In the data post-processing, we only included experiments with > 65% of samples above limit of detection. The limit of detection is set at 3 standard deviation above negative control values. Statistics run were one-way ANOVA with multiple comparison to one another and Dunnett’s post-hoc correction. * p<0.05; ** p<0.01. TNNI3 (cardiac troponin I), EPO (Erythropoietine), HPGDS (Hematopoietic prostaglandin D synthase), CA14 (Carbonic anhydrase 14), FES (FES proto-oncogene, Tyrosine Kinase)

### Summary

The outcomes of this paper suggest that chronic drug exposure in this MPS format elicits arrhythmic outcomes similar to those observed in published clinical trials[17, 22, 23, 35, 36, 38]. Specifically, the known arrhythmia risk of HCQ and AZM alone was recapitulated by our *in vitro* observation of APD_80_ increase in combination with arrhythmic events. Combination therapy also exhibited an increase in QT, but compared to monotherapy, was benign at inducing arrhythmogenic behaviors. This also corresponds with recent clinical findings [38]. Together, these data suggest that this our high content *in vitro* heart muscle model can aid clinicians in clinical trial design, rapidly predict the cardiac outcomes of polytherapy for SARS-CoV-2 treatment, and help to identify relevant biomarkers to monitor during clinical trials for potential COVID-19 therapeutics.

### Limitations

Clinical QT interval values represent the summation of all the electrical activity in the ventricles. We used APD_80_ as a proxy for clinical QT prolongation, which is a credible and common approach to compare directional effects and provide some mechanistic insight, but is not sufficiently sensitive for precise prediction of instability thresholds [30]. COVID-19 patients with drug-induced QT interval changes of > 60 ms or QT interval values above 500ms are considered high risk, and treatment is suspended[32]. Our cardiac muscle exhibited unphysiologically high APD_80_ values in response to chronic drug exposure. The mechanism for these large responses is unclear, but likely related to a combination of the hiPSC source, well-known modest maturity of hiPSC-CMs relative to the adult human heart, and possibly due to the altered current source-sink relationship in these very small tissues. Additionally, we have used a single patient line to perform this study, albeit with a significant number of replicates. By screening more patient lines one can achieve a clinically relevant dataset; although anticipated patient variability will require further expansion of the data size. In future work, this study can be extended to diseased cell lines to better understand the arrhythmic risk of patients with cardiovascular complications or comorbidities (i.e., diabetes) who are most likely to be seriously affected by SARS-CoV2.

### Study highlight

#### What is the current knowledge on the topic?

As the global pandemic of COVID-19 has expanded, clinicians were pressed to treat patients with new drug combinations, in the absence of regulatory approval. Even ten months after the first cases, FDA-approved medicines for the treatment of COVID-19 are just emerging with mixed results. There is a need for tools to rapidly screen for cardiac liability associated with potential therapeutics.

#### What question did this study address?

Does Hydroxychloroquine and Azithromycin COVID-19 polytherapy show synergetic cardiac liability when compared to their monotherapy?

#### What does this study add to our knowledge?

In this in vitro safety ‘clinical trial’, hydroxychloroquine or azithromycin alone showed significant APD80 (proxy for QT) prolongation and arrhythmia, whereas their combination polytherapy rescued instances of arrhythmias while increasing APD80.

#### How might this change clinical pharmacology or translational science?

This study demonstrates that a complex *in vitro* tissue model (cardiac MPS) can predict arrhythmias and rhythm instabilities under experimental conditions mimicking safety clinical trials. We also identified biomarkers associated with cardiac injury, which can be used to design clinical trial monitoring protocols.

## Acknowledgements

We thank Bruce Conklin (Gladstone Institutes, San Francisco, USA) for technical advice on the WTC iPSC line. We thank the Marvel Nanofabrication laboratory (UC Berkeley) and their staff for assistance and technical advices for microfabrication procedures.

## Author Contributions

All authors participated in the study design, analysis of the data, interpretation of the results and review of the manuscript; BC and VC conducted the main experiments and qualitative EAD characterization; BS helped with cell culture and MPS preparation; AGE helped with interpretation of electrophysiological recordings. HF developed the code to analyze action potential waveforms and Poincare plots; EM provided BeRST-1; BC, VC, AGE and KEH wrote the manuscript; BC prepared the figures and caption; KEH funded the work.

## Materials and Methods

### Cardiomyocyte Differentiation

The cardiomyocytes (CM) used in this study were derived from human induced pluripotent stem cells (hiPSC) via small molecular manipulation of the Wnt/ß-catenin signaling pathway [52] and were purified using glucose deprivation[53] to select for cardiomyocytes only.

### Fabrication and cell loading of Cardiac MPS

The microfluidic design for each MPS consisted of 4 identical cell culture chambers (1300 by 130 µm) with media channels running parallel on either side of the cell culture chambers. The microfluidic devices were fabricated from Polydimethylsiloxane (PDMS) using classic replica molding techniques [24, 29]. Upon loading, 4000 lactate purified hiPSC-CM were injected into each tissue chamber. The following day and every other day from then on, media was changed to our in-house ‘Maturation Media’ (MM) as described in [29]. MPS tissues were allowed to mature for at least 10 days before any subsequent experiments were performed.

### Drug Preparation For Pharmacology Studies

We based our study on the following protocols: Clinical drug administration for HCQ is 400 mg twice per day followed by 200 mg twice per day for 4 days. AZM is 500mg on day 1 followed by 250mg per day for the following 4 days. Based on clinical peak plasma concentration (C_max_) and area under curve for 24h[44][45], we exposed the MPS tissues to 0.24 μM HCQ at day 1 and 0.12 μM HCQ from day 2 to day 10; and to 2.67μg*h/ml / 24h = 0.111 μg/ml = 0.15 μM AZM at day 1 and 0.056 μg/ml (0.075 μM) AZM at day 2 to day 10.

### Thorough Action Potential Analysis As A Proxy For Clinical QT Interval Study And Arrhythmia Prediction

Clinically, drug-induced QT prolongation is a strong predictor of arrhythmic cardiotoxicity in patients. At the cellular level, AP prolongation and increased AP triangulation indicate slowed repolarization and are strong markers of whole heart QT prolongation and arrhythmia [37], translated by observations of Early afterdepolarizations (EADs) [46–49], and subsequent Torsade de Pointes[46]. Large beat-to-beat variation in AP duration is a specific indicator of repolarization instability, which can be readily visualized by Poincare plots, plotting CaD_80_ of each (n^th^) beat in the 30 second calcium recording, against CaD_80_ of the preceding beat (n-1)^th^, normalized to the CaD_80_ mean (**Figure 3A-C**). We also performed a qualitative and non-parametric evaluation of drug arrhythmogenesis by categorizing arrhythmic behaviors present in the calcium time-series (**Figure 3D-F**).

### Plasma Protein Profiling Using Olink Multiplex Panel

Effluents were sent to Olink Proteomics for quantification of proteins associated with toxicity and tissue damage. Olink Proteomics uses multiplex proximity extension assay (PEA) panels [50]. In this study, we have used the Organ Damage panel which consist of 92 unique markers of toxicity and cellular damage.

## Statistics

All statistics were calculated using GraphPad Prism. All electrophysiology data were analyzed with one way ANOVA repeated measures and Dunnett’s post-hoc correction with multiple comparison to day 0 and to one another was run. If some values were missing, mixed-effects model was run. Non parametric Chi-squared approximation was run for qualitative arrhythmic events assessment in a pairwise manner. Significance was determined with *p*-value < 0.05.

More detailed information can be found in the supplementary information.

