## Supplemental material for "In Vitro Safety “Clinical Trial” of the Cardiac Liability of Hydroxychloroquine and Azithromycin as COVID19 Polytherapy"

**Supplemental Materials and Methods**

**Cell Sourcing**
The human hiPSC line Wild Type C (WTC) harboring a single-copy of CAG-driven GCaMP6f knocked into the first Exon of the AAVS1 “safe harbor” locus [1] was used for all experiments in this study. It is available from the Coriell Repository (# GM25256 hiPSC from Fibroblast). More detailed information can be found in [1]*.*

**Cardiomyocyte Differentiation**
HiPSC stocks were thawed and plated on Matrigel hESC-Qualified Matrix (Corning; Corning, NY) coated tissue culture plates in mTeSR-1 (mTeSR-1; StemCell Technologies, Vancouver, Canada) containing 10µM of the Rho kinase inhibitor (RI) Y27632-dihydrochloride (Peprotech; Rocky Hill, NJ). Media was exchanged the next day to remove the inhibitor, and hiPSC were split three times at a density of 20,000 cells/cm2 to allow for recovery from cryopreservation. After three splits, hiPSC were seeded differentiation plates at 30-40,000 cells/cm2 to allow for full confluency. Initial seeding was considered as ‘day-3’ of differentiation. At day 0, when hiPSC were >90% confluent, differentiation was started following the protocol developed by Palecek et al. [[52](https://mail.google.com/mail/u/1/#m_7578038894609244351__ENREF_52)], which exposes the cells to 8µM CHIR99021 (Peprotech) in Roswell Park Memorial Institute Medium 1640 (RPMI) containing B-27 supplement without insulin (RPMI-I). At day 1, only RPMI-I was added to the cells. On day 2, cells were exposed to 5µM IWP-4 (Peprotech) in RPMI-I supplemented with 150µg/mL L-ascorbic acid (LAA) for two days. On day 4, only RPMI-I was given to the cells for another two days. From then onwards, RPMI containing standard B-27 supplement with insulin (RPMI-C) was added every second day. Once spontaneous contractions were observed around day 7 or 8 of differentiation, hiPSC-CM were washed several times with dPBS -Ca/Mg and left to soak for 10 minutes to disrupt cadherins. Cells were then exposed to 0.25% Trypsin (Thermo Fisher) for 10 minutes at 37C, and were pipetted gently to break up tissue into single cells and collected. Cells were pelleted at 300g for 5 minutes before being plated at a density of 100,000 cells/cm2 onto Matrigel, in RPMI-C with 10µM RI (‘day 0’ after replating). At day 1 after replating, medium was exchanged for RPMI-C. At day 3, cells were washed with dPBS to remove glucose and were exposed to RPMI 1640 (no glucose, no pyruvate, supplemented with 23mM sodium bicarbonate and 5mM Sodium L­-lacate[[53](https://mail.google.com/mail/u/1/#m_7578038894609244351__ENREF_53)]) for four to five days (exchanged every other day) to select for cardiomyocytes only. At day 8, cells were washed with dPBS and exposed to RPMI-C for two to three days to allow for recovery. Cardiomyocyte purity was characterized by flow cytometry with Cardiac Troponin T both before and after this procedure.

**Fabrication of Cardiac MPS**
The microfluidic design for each MPS consisted of 4 identical cell culture chambers (1300 by 130 µm) with media channels running parallel on either side of the cell culture chambers. All 8 media channels were connected to a single media inlet port and a single media outlet port (corresponding to the tubing in **Figure 1A**). Each cell culture chamber had its own cell loading port. Media channels and cell culture chambers were 60µm tall. An array of fenestrations (2µm x 2µm height/width and 40µm length) linked media channels to cell chambers, allowing for media change while protecting the tissue from shear stress [24] (**Figure 1B**).
The microfluidic devices were fabricated from Polydimethylsiloxane (PDMS) using classic replica molding techniques. The basic process is described in our previous work [2, 3] and has been optimized. A master mold featuring the negative structures of the fluidic channels was prepared in a 2-step photolithography process. First the 2µm thick fenestration layer was patterned on a clean and water-free silicon wafer by spin-coating SU8 2002 (MicroChem) in a 2-step process (500rpm, 10sec, 100rpm/sec acceleration; 2000rpm, 30sec, 300rpm/sec acceleration). A soft bake of 1min at 95C was performed prior to 80mJ/cm^2^ exposure. The post exposure bake was 2min at 95C, followed by a development in SU8 developer (30sec on shaker, 15sec on sonicator and 30sec on shaker). For the second layer SU8 3050 (MicroChem) was spun to 60µm thickness (500rpm, 10sec, 100rpm/sec acceleration; 3000rpm, 30sec, 300rpm/sec acceleration). The soft bake was performed for 10min at 65C followed by 45min at 95C with very slow up and down ramping of the temperature. Alignment markers were used to position the 2^nd^ layer mask over the fenestration layer features during the exposure step (160mJ/cm^2^ ). The post exposure bake was 2min at 65C and 10min at 95C again with slow ramping. SU8 developer was applied on a shaker for 20min and sonicator for 15min. A final hard baked was performed at 180C for 30-45min.
The cardiac MPS was formed by replica molding from the silicon wafer with Polydimethylsiloxane (PDMS; Sylgard 184 kit, Dow Chemical, Midland, MI) at a 10:1 ratio of Sylgard base to crosslinker. The inlets and outlets were punched into the PDMS with a 0.75 µm biopsy punch before bonding to glass slides using oxygen plasma (PETS inc. RIE system) with the following protocol: RF power 21W, 24sec, 100% oxygen flow (600mTorr).

**Self-Assembly of Cardiac Microtissues within Cardiac MPS**
Lactate purified hiPSC-CM were singularized with 0.25% trypsin for 5 min and suspended to a density of 1.33x10^6^ cells/mL in EB20 media (Knock Out DMEM (Gibco, 10829-01820%) with 20% FBS (Gibco, 16000-0), 1% MEM non-essential amino acids (Gibco, 11140-050), 1% Glutamax (Gibco, 35050-061) and 400nM beta-mercaptoethanol (Gibco, 21985023)) freshly supplemented with 10µM ROCK-inhibitor Y27632 (Sigma, SCM075). 3µL of the cell suspension, corresponding to 4000 cells, was injected into the cell loading port of each tissue chamber. To move the cells into the cell chamber MPS were centrifuged twice at 300g for 3 minutes: first in horizontal and then in vertical position. The loading tips were then sealed. Chambers that were not filled at this point were discarded. MPS were incubated at 37C for 1-2h to allow formation of a dense cell cluster and avoid backflow of cells when adding media. EB20 media supplemented with 10µM Y27632 (200uL per MPS) was added to the media inlet of each MPS and flow was initiated by applying gentle vacuum at the media outlet. The following day and every other day from then on, media was changed to our in-house ‘Maturation Media’ (MM) as described in [3]. MPS tissues were allowed to mature for at least 10 days before any subsequent experiments were performed. When tissues were fed (every 2 days for maintenance or every day for drug studies), 200μL of media was added to the inlet tip and media would flow gravimetrically to the outlet until equilibrium was reached.

**Preparation of Maturation Media (MM)**
A base media was prepared from RPMI 1640 powder (Sigma, R1383-10X1L) and supplemented with 0.5g/L D-glucose (Fisher, BP350-1), 10mM D-galactose (Sigma, G5388) and 2g/L sodium bicarbonate (Fisher, S233-500) . A 9% bovine serum albumin (BSA; Fisher, BP1605-100) solution was prepared in base media and adjusted to pH 7.4. Aliquots of fatty acid (FA) stocks were prepared from oleic acid (OA ; Sigma, O1383-5G) and palmitic acid (PA; Sigma, P0500-10G ; 100mg/ml in DMSO) and stored at -20C. OA was prediluted 1:10 in DMSO. PA and OA were added to the 9% BSA solution at 10.23ug/ml for PA and 0.8mM OA. To dissolve the FA the solution was heated to 60C and sonicated for 5min. The final MM was prepared by mixing 1 part FA solution and 3 parts MM base media yielding a final concentration of 2.5575 ug/ml PA and 0.2 mM OA. Finally, the media was supplemented with 2% B27 (Gibco, 17504-044) and 150μg/ml ascorbic acid (Fisher, AC105021000) and sterile filtered.

**Drug preparation for pharmacology studies**
We performed all the pharmacology experiments in MM with 5nM BeRST-1 dye to guarantee a high-quality signal. On the day of pharmacology study, drugs were weighed and dissolved in an adequate solvent to create a stock solution (**Supp. Figure 1E**). From that stock, the highest test dose was prepared and serial dilutions were made for lower doses. Vehicle concentration was kept the same in all doses including dose 0 and controls.

In more detail, hydroxychloroquine sulfate (PHR1782-1G, Sigma) was dissolved in PBS (without calcium or magnesium) at 4.34mg/ml and sterile filtered to create a 10mM stock solution. The stock was always prepared fresh on the day of the experiment. Azithromycine (75199-25MG-F, Sigma) was dissolved in 100% ethanol to create a 10 mg/ml (=13.35mM) stock. The stock was stored at -20°C for up to 3 days. The chronic drug exposure doses were chosen to mimick clinical trials (HCQ[4], AZM [5]) based on patients’ serum concentration. While there is some variability in clinical protocols, we based our study on the following commonly used protocols: Clinical drug administration for HCQ is 400 mg twice per day followed by 200 mg twice per day for 4 days. For 200mg, the clinical peak plasma concentration (C_max_) for oral HCQ sulfate is 50.3 ng/mL (0.12 μM) at t_max_ of 3.74 h after administration[6]. Therefore, to mimick clinical trials, the MPS tissues were exposed to 0.24 μM HCQ at day 1 and 0.12 μM HCQ from day 2 to day 10. Clinical administration of AZM is 500mg on day 1 followed by 250mg per day for the following 4 days. Following 500mg administration, the area under curve for 24h is 2.67 (+/-0.92) μg*h/ml and C_max_ 0.405 (+/-0.161) μg/ml [7]. Therefore, to mimick clinical trials, the MPS tissues were exposed to 2.67μg*h/ml / 24h = 0.111 μg/ml = 0.15 μM AZM at day 1 and 0.056 μg/ml (0.075 μM) AZM at day 2 and following. We continued AZM application for all 10 days of the study.

**Experimental Setup**
24h prior to any experiment, MPS tissues were stained with the action potential dye BeRST-1 (500nM), synthesized and QC as described my Miller et al. (2018). During imaging, MPS were kept at 37°C on a heated microscope stage (Tokai Hit, Gendoji-cho, Japan). MPS were placed onto the heat plate for 20 mins to let them stabilize before imaging.
Initial recordings were performed on day 0 prior to any drug exposure. After recordings, supernatants from the inlet and outlet were collected and frozen at -80C for proteomics analysis. Drug doses were always prepared fresh directly before use. Drug containing media (200uL per MPS) was applied daily via the media inlet port and incubated for 24h in a cell culture incubator at 37C where gravimetric perfusion occurred as the liquid levels in inlet and outlet tip equilibrated. The same steps were repeated for 10 days. The same MPS tissues were repeatedly recorded daily for 10 days. Drug doses are outlined in section “Drug preparation for pharmacology studies”.
Control tissues were handled equivalent to tissues with drug exposure : every day fresh media containing a solvent concentration identical to the drug vials was applied. Control MPS were subjected to the same imaging protocol, and their effluents were also collected.

**Image Acquisition for Pharmacology Studies**
Videos of calcium (GCaMP) and voltage (BeRST-1) epifluorescence, were recorded daily (every 24h) in spontaneously beating tissues, and directly before administration of a fresh drug dose. NIKON TE300HEM microscope with a HAMAMATSU digital CMOS camera C11440 / ORCA-Flash 4.0 was used and enabled 100 frames per second acquisition rate. For fluorescence imaging, a Lumencor SpectraX Light Engine and filtered with a QUAD filter (Semrock) was used. GCaMP videos (6s and 30s long) used the Cyan LED, 470nm (4x4 binning, 10ms), while BeRST-1 videos (6s long) used the Far-red LED, 640ms (4x4 binning, 10ms). Videos were acquired using Nikons Nikon NIS-Elements software.
Post-experiment processing was performed with an in-house python library. This library performs automated background subtraction and normalization, and eliminates baseline drift due to bleaching by fitting a polynomial to the baseline via half-quadratic minimization[8]. The resulting traces were used for quantitative analysis of the action potential by calculating metrics such as 80% and 30% action potential duration (APD_80_ and APD_30_), triangulation ((APD_80_-APD_30_)/ APD_80_) and beat rate. These analyses were also automated by the same python library.

**Thorough Action Potential Analysis As A Proxy For Clinical QT Interval Study And Arrhythmia Prediction**

Clinically, the QT interval describes the period between Purkinje activation (Q) and ventricular repolarization (T), and drug-induced QT prolongation is a strong predictor of subsequent arrhythmic cardiotoxicity in patients (**Figure 1D**, top). Torsades de pointes (TdP) is a particularly dangerous form of polymorphic ventricular tachycardia and the most common arrhythmic manifestation of drugs that induce dangerous QT prolongation. TdP is a ventricular tachycardia triggered by EADs, where the amplitude and configuration of ECG measurements vary continously. At the cellular level, AP prolongation and increased AP triangulation indicate slowed repolarization and are strong markers of whole heart QT prolongation and arrhythmia [9]. Early afterdepolarizations (EADs) represent a transition from slowed cellular repolarization to unstable (chaotic) repolarisation[10-13] at the cellular level, and are a harbinger of whole-heart TdP[10]. These characteristics formed the basis of our analysis of the MPS voltage (BeRST-1) and calcium (GCaMP) recordings, which were assessed for drug-induced changes to both stable beats (e.g. APD_30_, APD_80_ etc.) and unstable cellular behaviors (e.g. EADs etc.).

**Correlation between APD_80_ and CaD_80_**

Analysis of the correlation between calcium transient duration (CaD80) and action potential duration (APD80) shows a strong linear correlation (Y = 0.8163*X + 109.1) with R2=0.86 in the relevant range of APD80=250 to 800ms and slightly higher variation above 800ms (**Supp.Figure 1D**). This good correlation confirms that CaD80 values can be used in place of APD80 values to predict drug induced changes of the beat length.

**Stable beating analyses:**

Since patients with QT interval > 500ms are considered at risk for TdP [14], all tissues with baseline APD_80_ > 500ms were *a priori* excluded from the study. Similarly, because APD is inversely proportional to beating frequency [15] (**Figure 1D**, bottom), APD measures were first corrected for differences in beating frequency via the Fridericia method[16]. As per Hondeghem et al., [9] if APD prolongation is accompanied by instability or triangulation then EADs and arrhythmia are very likely to follow. For these reasons we used the drug-induced changes to APD_80_, and AP triangulation as primary measures of drug arrhythmogenicity in recordings that exhbited stable AP waveforms.

**Analyses of unstable beating:**
Large beat-to-beat variation in AP duration is a specific indicator of repolarization instability. EADs represent a form of dynamical chaos in repolarization, and APD alternans is a period-2 dynamic repolarization instability. Both behaviors can be readily visualized by Poincare plots (**Figures 3A-C**). These were again generated by an in-house python script plotting CaD_80_ of each (n^th^) beat in the 30 second calcium recording, against CaD_80_ of the preceding beat (n-1)^th^, normalized to the CaD_80_ mean. Identical CaD_80_ values in sequence appear as a single point, consistent CaD increase or decrease (anti-arrhythmic) will cluster around the center of the graph and if large deviations between successive CaD_80s_ appear (pro-arrhythmic), points will deviate from the center and meander, giving rise to disorganized polygons[9]. We also performed a qualitative and non-parametric evaluation of drug arrhythmogenesis by categorizing arrhythmic behaviors present in the calcium time-series (spontaneous 30 second recordings, **Figure 3D-F**). The arrhythmic categories included EADs, DADs, irregular beating (successive beats with different APD or resting time), TdP (continous changes in amplitude), overt changes to automaticity (as suggested by Blinova et al. [15]), and weak or quiescent tissues (suggesting loss of resting membrane polarization).

**Plasma Protein Profiling Using Olink Multiplex Panel**
Effluents were stored at -80C and all samples were sent together on dry ice to Olink Proteomics for quantification of proteins associated with toxicity and tissue damage. Olink Proteomics uses multiplex proximity extension assay (PEA) panels [17]. The basis of PEA is a dual-recognition immunoassay, where two matched antibodies labelled with unique DNA oligonucleotides simultaneously bind to a target protein in solution. This brings the two antibodies into proximity, allowing their DNA oligonucleotides to hybridize, serving as template for a DNA polymerase-dependent extension step. This creates a double-stranded DNA “barcode” which is unique for the specific antigen and quantitatively proportional to the initial concentration of target protein. The hybridization and extension are immediately followed by PCR amplification and the amplicon is then finally quantified by microfluidic qPCR using Fluidigm BioMark HD system (Fluidigm Corporation. South San Francisco, California). In this study, we have used the Organ Damage panel which consist of 92 unique markers of toxicity and cellular damage (https://www.olink.com/products/organ-damage-panel/). In the data post processing, we only included biomarkers with > 65% of samples above limit of detection. The limit of detection is set at 3 standard deviations above negative control values, making it quite conservative.

**Statistics**
All statistics were calculated using GraphPad Prism. All electrophysiology data were analyzed with one way ANOVA repeated measures and Dunnett’s post-hoc correction with multiple comparison to day 0 and to one another was run. If some values were missing, mixed-effects model was run. Non parametric Chi-squared approximation was run for qualitative arrhythmic events assessment in a pairwise manner. Significance was determined with *p*-value < 0.05.

**Supplemental Figure Legends**

**Supplementary Figure 1. Triangulation analysis of chronic either hydroxychloroquine (HCQ), azithromycin (AZM), or their polytherapy.** Spontaneous triangulation values from action potential traces in tissues treated with HCQ (**A**), AZM (**B**) and polytherapy (**C**). For each tissue, triangulation is defined as the difference between APD_80_ to APD_30_, normalized to APD_80_. (**D**) The solid lines shows the linear correlation of CaD80 and APD80 values (Y = 0.82*X + 109; R^2^=0.86) and the the dashed lines indicate the 90%-confidence intervals.

**Supplemental Tables**

**Supplementary Table 1.** Table describing drug references, primary purpose, clinical C_max_ and doses used in this study.

| **Drug  (Reference)** | **Primary Purpose** | **Clinical C_max_** | **Chronic drug dose**  (based on clinical trials area under curve over 24h exposure) |
| --- | --- | --- | --- |
| Hydroxychloroquine - HCQ *(PHR1782-1G, Sigma)* | Anti-malarial | 1μM | **Day1** = 0.24μM  **Day2-10** = 0.12μM |
| Azithromycin - AZM *(75199-25MG-F, Sigma)* | Macrolide Antibiotic | 0.67μM | **Day1** = 0.15 μM  **Day2-10** = 0.075 μM |
| HCQ+AZM combination | SARS-CoV2 |  | same as each respective concentration but combined |
