## Supplementary figures and images for "In Vitro Safety “Clinical Trial” of the Cardiac Liability of Hydroxychloroquine and Azithromycin as COVID19 Polytherapy"

### Supplemental figure

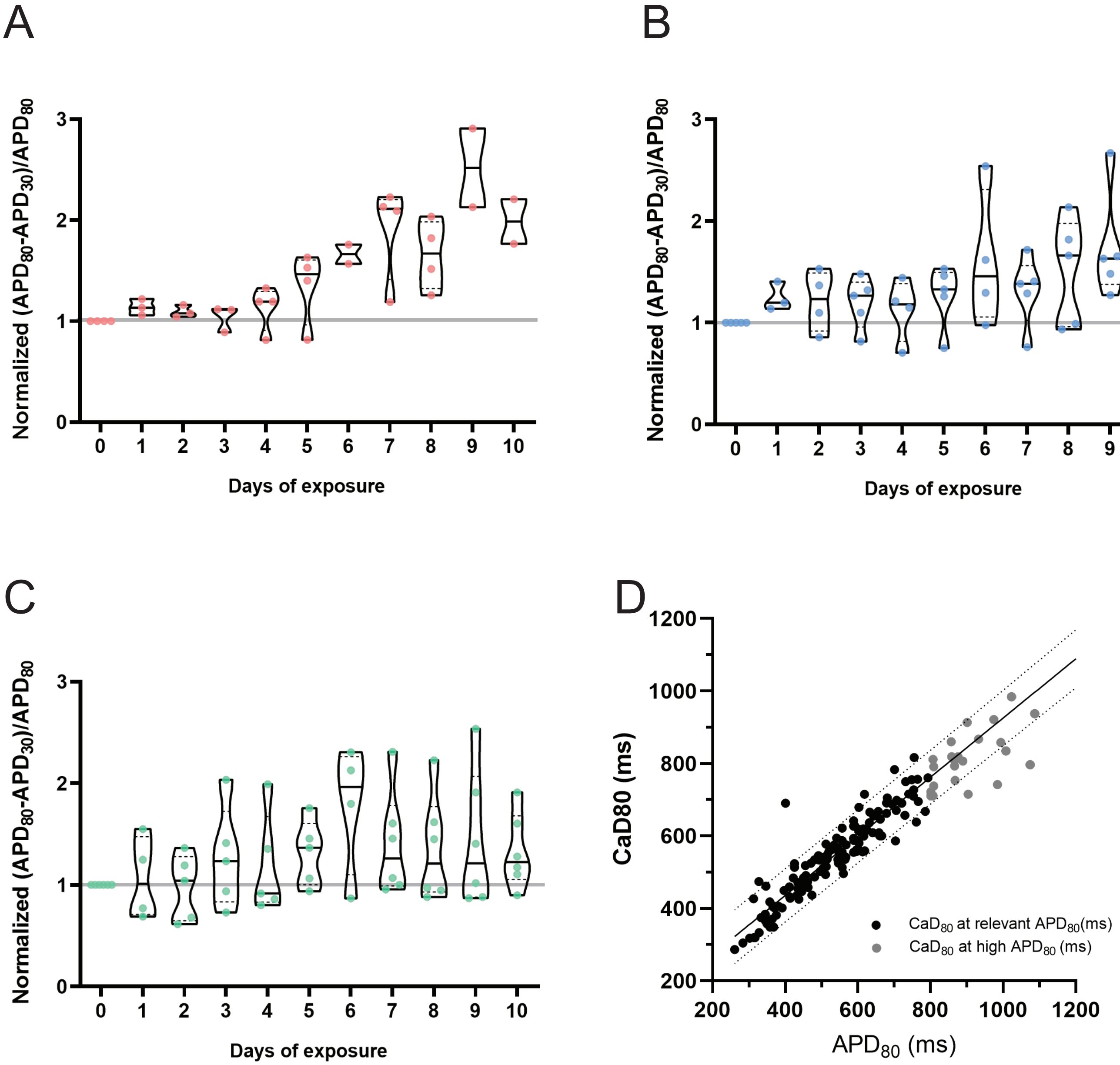
